# *Salmonella* genomic island 3 is an integrative and conjugative element and contributes to copper and arsenic resistance of *Salmonella enterica*

**DOI:** 10.1101/564534

**Authors:** Nobuo Arai, Tsuyoshi Sekizuka, Yukino Tamamura, Masahiro Kusumoto, Atsushi Hinenoya, Shinji Yamasaki, Taketoshi Iwata, Ayako Watanabe-Yanai, Makoto Kuroda, Masato Akiba

## Abstract

*Salmonella* genomic island 3 (SGI3) was first described as a chromosomal island in *Salmonella* 4,[5],12:i:-, a monophasic variant of *Salmonella enterica* subsp. *enterica* serovar Typhimurium. The SGI3 DNA sequence detected from *Salmonella* 4,[5],12:i:-isolated in Japan was identical to that of a previously reported one across entire length of 81 kb. SGI3 consists of 86 open reading frames, including a copper homeostasis and silver resistance island (CHASRI) and an arsenic resistance operon in addition to genes related to conjugative transfer and DNA replication or partitioning, suggesting that the island is a mobile genetic element. We successfully selected transconjugants that acquired SGI3 after filter mating experiments using the *S. enterica* serovars Typhimurium, Heidelberg, Hadar, Newport, Cerro, and Thompson as recipients. Southern blot analysis using I-CeuI-digested genomic DNA demonstrated that SGI3 was integrated into a chromosomal fragment of the transconjugants. PCR and sequencing analysis demonstrated that SGI3 was inserted into the 3′ end of the tRNA genes *pheV* or *pheR*. The length of the target site was 52 or 55 bp, and a 55-bp *attI* sequence indicating generation of the circular form of SGI3 was also detected. The transconjugants had a higher MIC against CuSO_4_ compared with the recipient strains under anaerobic conditions. Resistance was defined by the *cus* gene cluster in the CHASRI. The transconjugants also had distinctly higher MICs against Na_2_HAsO_4_ compared with recipient strains under aerobic conditions. These findings clearly demonstrate that SGI3 is an integrative and conjugative element and contributes to the copper and arsenic resistance of *S. enterica*.

## INTRODUCTION

*Salmonella enterica* subsp. *enterica* serovar Typhimurium (*Salmonella* Typhimurium) and its monophasic variant (*Salmonella* 4,[5],12:i:-) are major causes of gastroenteritis in humans and animals worldwide (1-4). In particular, *Salmonella* 4,[5],12:i:-infection of humans and animals has increased considerably in European countries since the mid-1990s. The epidemic *Salmonella* 4,[5],12:i:-strain is called the ‘European clone’ and is characterized by sequence type 34 (ST34) and possession of a composite transposon containing antimicrobial resistance genes and a chromosomal island, *Salmonella* genomic island 3 (SGI3). This clone has been spread from Europe to the other area, including North America and Asia (1, 2, 5-9).

In Japan, bovine salmonellosis caused by *Salmonella* 4,[5],12:i:-has been increasing in the last decade (6, 10, 11). We performed a phylogenetic analysis using whole-genome sequencing data of 119 *Salmonella* Typhimurium and *Salmonella* 4,[5],12:i:-isolates obtained in Japan and Italy and identified 9 distinct clades as genotypes for epidemiological analysis. Then, we established an allele-specific PCR-based genotyping system detecting a clade-specific single nucleotide polymorphism (SNP) to rapidly identify the clade of each isolate. Clade 9 was the most prevalent in recent years among 955 *Salmonella* Typhimurium and *Salmonella* 4,[5],12:i:-isolates obtained from food animals in Japan between 1976 and 2017. This clade mainly consists of *Salmonella* 4,[5],12:i:- and shares the same characteristics as the European clone. We speculate that the increased prevalence of clade 9 isolates in Japan is a part of the pandemic of the European clone (12).

All the clade 9 isolates harbored a composite transposon containing antimicrobial resistance genes defining resistance to ampicillin, streptomycin, sulfonamide, and tetracycline. However, these antimicrobials do not appear to be a main selection pressure because some former epidemic clones of *Salmonella* Typhimurium showed similar resistance patterns (13, 14). Clade 9 isolates also harbored SGI3, which contains genes for resistance to copper, zinc, and arsenic, and the European clone exhibits an increased minimal inhibitory concentration (MIC) for copper sulfate in rich broth culture (15). As copper and zinc are generally added to feed of livestock to achieve growth promotion and increased feed efficiency (16), these feed additives might be an important selection pressure for this clone.

Integrative and conjugative elements (ICEs) are self-transmissible between chromosomes of different cells (17, 18) and are widely distributed among both Gram-negative and Gram-positive bacteria (19). ICEs include genes defining a type IV secretion system, which contributes to conjugative transfer, in addition to an integrase and an excisionase (17, 19). ICEs can excise themselves from the donor chromosome, form a circular intermediate, transfer by conjugation and reintegrate into the recipient chromosome. Each ICE has variety of cargo genes that contribute to antimicrobial resistance (20-24), biofilm formation (25), metabolism of alternative carbon sources (26), and degradation of the persistent chemical substances (27, 28). Hence, horizontal gene transfer by ICEs plays an important role for bacterial evolution.

SGI3 includes genes encoding the type IV secretion system and a site-specific recombinase, suggesting that SGI3 might be an ICE and provide heavy metal resistance to the host. However, functions of SGI3 as an ICEs are unproved to date. In this study, we determined the transfer frequencies of SGI3 using clade 9 strains as donors and other *Salmonella* strains as recipients, and the susceptibility of the transconjugants to heavy metals was compared with that of their parental strains to prove that SGI3 was a functional ICE. Moreover, we constructed deletion mutants of the copper resistance gene clusters among SGI3 and compared the susceptibility to copper with that of the parental strains to identify the functional gene clusters that contributed to copper resistance.

## MATERIALS AND METHODS

### Bacterial strains

The strains used in this study are listed in Table 1. *Salmonella* Typhimurium LT2 (29) was purchased from American Type Culture Collection (Manassas, VA, USA). Wild-type strains of *S. enterica* were obtained from cattle, swine, or birds in Japan between 1997 and 2011. These strains were isolated by the staff of the local animal hygiene service centers. *Salmonella* spp. were identified based on colony morphology on selective media and biochemical properties, as previously described (30). Serovar identification was performed by microtiter and slide agglutination methods according to the White-Kaufmann-Le Minor scheme (31). Whole genome raw sequence reads of *S. enterica* strains L-3785, L-3838, L-3841, L-4125, L-4131, L-4153, and L-4239 were determined and deposited in the DDBJ Sequence Read Archive in the previous study (12). *Salmonella* 4,[5],12:i:-L-3841, which harbors SGI3 in its chromosome, was used as a donor in conjugation experiment. All strains but L-3838 and L-3841 were used as the recipients in conjugation experiments. Rifampicin-resistant mutants of all recipients were selected on DHL agar plates (Nissui Pharmaceutical Co., LTD., Tokyo, Japan) containing 100 µg/ml of rifampicin (FUJIFILM Wako Pure Chemical Corp., Osaka, Japan).

**Table 1.** Bacterial strains used in this study

| Strain | Serovar | Description | SNP genotype<sup>a</sup> | Accession number<sup>a</sup> | Reference or source |
| --- | --- | --- | --- | --- | --- |
| LT2 | Typhimurium | ATCC700726 | 3 | AE006468.2 | 29 |
| LT2TC*pheV* | Typhimurium | LT2 transconjugant which acquired SGI3 at *pheV* locus | 3 | NA | This study |
| LT2TC*pheR* | Typhimurium | LT2 transconjugant which acquired SGI3 at *pheR* locus | 3 | NA | This study |
| LT2TC*pheR* ΔSGI3 | Typhimurium | SGI3 deletion mutant derived from LT2TC | 3 | NA | This study |
| LT2TC*pheR* Δ*pco* | Typhimurium | *pco* gene cluster deletion mutant derived from LT2TC*pheR* | 3 | NA | This study |
| LT2TC*pheR* Δ*cus* | Typhimurium | *cus* gene cluster deletion mutant derived from LT2TC*pheR* | 3 | NA | This study |
| L-3838 | 4,[5],12:i:- | Wild type | 9 | AP019374 | This study |
| L-3841 | 4,[5],12:i:- | Wild type | 9 | AP019375 | This study |
| L-3841ΔSGI3 | 4,[5],12:i:- | SGI3 deletion mutant derived from L-3841 | 9 | NA | This study |
| L-3841Δ*pco* | 4,[5],12:i:- | *pco* gene cluster deletion mutant derived from L-3841 | 9 | NA | This study |
| L-3841Δ*cus* | 4,[5],12:i:- | *cus* gene cluster deletion mutant derived from L-3841 | 9 | NA | This study |
| L-3900 | Typhimurium | Wild type | 1 | NA | 10 |
| L-3900TC | Typhimurium | L-3900 transconjugant which acquired SGI3 at *pheV* or *pheR* locus | 1 | NA | This study |
| L-4131 | Typhimurium | Wild type | 2 | SAMD00097416 | 12 |
| L-4131TC | Typhimurium | L-4131 transconjugant which acquired SGI3 at *pheV* or *pheR* locus | 2 | NA | This study |
| L-3785 | 4,[5],12:i:- | Wild type | 4 | SAMD00097434 | 12 |
| L-3785TC | 4,[5],12:i:- | L-3785 transconjugant which acquired SGI3 at *pheV* or *pheR* locus | 4 | NA | This study |
| L-4153 | Typhimurium | Wild type | 5 | SAMD00097444 | 12 |
| L-4153TC | Typhimurium | L-4153 transconjugant which acquired SGI3 at *pheV* or *pheR* locus | 5 | NA | This study |
| L-4125 | Typhimurium | Wild type | 6 | SAMD00097448 | 12 |
| L-4125TC | Typhimurium | L-4125 transconjugant which acquired SGI3 at *pheV* or *pheR* locus | 6 | NA | This study |
| L-4239 | Typhimurium | Wild type | 7 | SAMD00097461 | 12 |
| L-4239TC | Typhimurium | L-4239 transconjugant which acquired SGI3 at *pheV* or *pheR* locus | 7 | NA | This study |
| L-3569 | Typhimurium | Wild type | 8 | NA | This study |
| L-3569TC | Typhimurium | L-3569 transconjugant which acquired SGI3 at *pheV* or *pheR* locus | 8 | NA | This study |
| L-2707 | Heidelberg | Wild type | NA | NA | This study |
| L-2707TC | Heidelberg | L-2707 transconjugant which acquired SGI3 at *pheV* or *pheR* locus | NA | NA | This study |
| L-2819 | Hadar | Wild type | NA | NA | This study |
| L-2819TC | Hadar | L-2819 transconjugant which acquired SGI3 at *pheV* or *pheR* locus | NA | NA | This study |
| L-3037 | Newport | Wild type | NA | NA | This study |
| L-3037TC | Newport | L-3037 transconjugant which acquired SGI3 at *pheV* or *pheR* locus | NA | NA | This study |
| L-3038 | Infantis | Wild type | NA | NA | This study |
| L-3046 | Dublin | Wild type | NA | NA | This study |
| L-3202 | Choleraesuis | Wild type | NA | NA | This study |
| L-3289 | Cerro | Wild type | NA | NA | This study |
| L-3289TC | Cerro | L-3289 transconjugant which acquired SGI3 at *pheV* or *pheR* locus | NA | NA | This study |
| L-3241 | Enteritidis | Wild type | NA | NA | This study |
| L-4000 | Thompson | Wild type | NA | NA | This study |
| L-4000TC | Thompson | L-4000 transconjugant which acquired SGI3 at *pheV* or *pheR* locus | NA | NA | This study |

<sup>a</sup>NA, not applicable

### Identification of SGI3 structure in wild-type strains

To determine the complete genome sequences of *Salmonella* 4,[5],12:i:-strains L-3838 and L-3841, high-molecular weight DNA was extracted followed by long-read sequencing using a PacBio Sequel sequencer (Sequel SMRT Cell 1M v2 [four/tray]; Sequel sequencing kit v2.1; insert size, approximately 10 kb). The long-read libraries were prepared with a SMRTbell library using a SMRTbell template prep kit 1.0 (PacBio, Menlo Park, CA, USA) with barcoding adapters according to the manufacturer’s instructions. These sequencing data were produced with more than 50-fold coverage, and pre-assembled reads were generated using SMRT Link software v5. *De novo* assembly with pre-assembled reads was performed using Canu version 1.4 [PMID: 28298431], Minimap version 0.2-r124 [PMID: 27153593], racon version 1.1.0 [PMID: 28100585], and Circlator version 1.5.3 [PMID: 26714481]. Error correction of assembled sequences was performed using Pilon version 1.18 [PMID: 25409509] with Illumina short reads previously deposited in the SRA database (12). Open reading frames (ORFs) were predicted and automatically annotated by DDBJ Fast Annotation and Submission Tool (DFAST) (32). Then, each ORF in SGI3 structure was confirmed using the Basic Local Alignment Search Tool (BLAST; https://blast.ncbi.nlm.nih.gov/Blast.cgi) in reference to SGI3 in *Salmonella* 4,[5],12:i:-strain TW-Stm6 (Accession number CP019649) (33).

### Pairwise alignment of SGI3 with relevant DNA sequences

Pairwise alignment of SGI3 and other mobile genetic elements was performed using the BLAST Search Tool followed by a homology search using the NUCmer alignment program from MUMmer in GENETYX^®^ version 13 (GENETYX Co., Ltd., Tokyo, Japan). The alignment image was visualized using Easyfig version 2.2.2 (34).

### Conjugation experiment

Conjugation experiments were performed using the filter mating method (35) with slight modifications. Briefly, donor and recipients were grown in Luria-Bertani (LB) broth (Becton, Dickinson and Company, Franklin Lakes, NJ) at 37°C for 18 hours. Aliquots (50 µl) of both cultures were transferred to 5 ml fresh LB broth and incubated at 37°C for 4 hours with shaking until exponential phase growth was achieved. Then, 0.5 ml of donor and 4.5 ml of recipient were mixed and trapped on a sterile mixed cellulose ester filter (0.45 µm pore size, Toyo Roshi Kaisha, Ltd. Tokyo, Japan). The filter was placed on a LB agar plate, which was incubated at 37°C for 20 hours. Bacterial cells were removed from filters by vortexing with 1 ml of sterile saline solution. Transconjugants were selected on DHL agar plates containing rifampicin (100 µg/ml) and Na_2_HAsO_4_ (8 mM) (FUJIFILM Wako Pure Chemical). Donor was selected on DHL agar plates with Na_2_HAsO_4_ (8 mM). Transfer frequency was calculated as the number of transconjugants divided by the number of donor cells. We repeated the conjugation experiments up to thrice for each recipient to obtain transconjugants.

### PCR

Primers used in this study are listed in Table S1. SNP genotyping was performed to identify the genetic background of *Salmonella* Typhimurium and *Salmonella* 4,[5],12:i:-using primer pairs SNP3_F/SNP3_R and SNP9_F/SNP9_R (12). IS*200* located between the genes *fliA* and *fliB* of *Salmonella* Typhimurium can be used as a specific marker of this serovar (36). The primer pair FFLIB/RFLIA was used for amplification of the intergenic region by PCR to determine whether the tested isolate is *Salmonella* Typhimurium. The amplicon sizes from *Salmonella* Typhimurium, *Salmonella* 4,[5],12:i:-, and other serovars were expected to be 1,000, 1,000, and 250 bp, respectively (37). To localize SGI3, we amplified the junction region between SGI3 and the recipient chromosome. Primer pairs SGI3_LJ_F/SGI3_LJ_R and SGI3_RJ_F/SGI3_RJ_R were used to amplify the left and right junction of SGI3 at the *pheR* locus, respectively. Primer pairs SGI3_LJ_F2/SGI3_LJ_R and SGI3_RJ_F/SGI3_RJ_R2 were used to amplify the left and right junction of SGI3 at *pheV* locus, respectively. To confirm the occurrence of the circular form of SGI3, the primer pair SGI3_RJ_F/SGI3_LJ_R was used. All of the oligonucleotides used in this study were purchased from Hokkaido System Science (Hokkaido, Japan).

A single colony (2 mm in diameter) was suspended in 50 µl of 25 mM NaOH. After boiling for 5 min, the suspension was neutralized by adding 4 µl of 1 M Tris-HCl (pH 8.0) and centrifuged at 20,000 × *g* for 5 min, and 1 µl of the supernatant was used as template DNA. To confirm the occurrence of the circular form of SGI3, template DNA was prepared by collecting bacterial cells after shaking in LB-broth at 37°C for 4 hours.

SNP genotyping was performed using previously described PCR conditions (12). To detect the junction region of SGI3 and the host chromosome, PCR was performed in a 20-µl reaction mixture containing template DNA, 0.3 µM each primer, 0.2 mM deoxynucleotide triphosphates (dNTPs), PCR buffer, and 0.5 U *TaKaRa Ex Taq*^®^ Hot Start Version (Takara Bio Inc., Shiga, Japan) using 25 amplification cycles of 95°C for 30 s, 58°C for 30 s, and 72°C for 60 s. To detect the junction region of circular form of SGI3, PCR was performed in the same reaction mixture using 30 amplification cycles of 95°C for 30 s, 60°C for 30 s, and 72°C for 50 s.

PCR products were separated on 2% agarose gel and visualized by staining with ethidium bromide. DNA sequences of each fragment were determined by Sanger DNA sequencing using a BigDye Terminator cycle sequencing kit and Applied Biosystems 3130 DNA analyzer (Thermo Fisher Scientific, Inc., Waltham, MA). Nucleotide sequence assembly was performed using the ATGC sequence assembler of GENETYX^®^ ver.13.

### Localization of SGI3 by Southern blot analysis

To confirm the chromosomal location of SGI3, pulsed-field gel electrophoresis (PFGE) followed by Southern blot analysis was performed as described previously (38) with slight modifications. Briefly, plugs for I-CeuI restriction were prepared from total DNA of five isolates, including L-3838, L-3841, LT2TC*pheV*, LT2TC*pheR*, and LT2, according to the PulseNet protocol (39) with slight modifications. The cell culture for plug preparation had an optical density at 600 nm of 2. The final concentration of proteinase K (FUJIFILM Wako Pure Chemical) in the plugs was 1 mg/ml, and samples were washed with 10 mM Tris-HCl containing 1 mM EDTA (TE) buffer four times. The plugs were digested with 20 U/plug of I-CeuI (New England BioLabs, Inc., Ipswich, MA) or 40 U/plug of XbaI (Takara Bio Inc.) at 37°C for 6 hours. XbaI-digested DNA from *Salmonella* Braenderup str. HB9812 was used as a molecular standard marker. The digested fragments were separated by PFGE (6.0 V/cm for 21 hours with a pulsing time linearly ramped from 2.2s to 63.8 s) using the CHEF DR-II apparatus (Bio-Rad Laboratories, Inc., Hercules, CA) on a 1% agarose gel. The separated I-CeuI fingerprint was transferred to positively charged nylon membranes (Roche Diagnostics Co., Indianapolis, IN) and hybridized with PCR-generated digoxigenin-labeled probes (Roche Diagnostics) using DIG Easy Hyb hybridization solution (Roche Diagnostics) according to the manufacturer’s instructions. Primer pairs 23S-For/23S-Rev, pcoA-F/pcoA-R, and int-F/int-R were used to generate the probes for the detection of 23S rRNA, *pcoA*, and integrase genes, respectively (Table S1).

### Construction of gene replacement vectors and generation of deletion mutants

To generate deletion mutants of SGI3, *pco* gene cluster, and *cus* gene cluster, gene replacement was performed as described previously (40) with slight modifications. For SGI3, upstream and downstream regions of SGI3 were amplified and connected by three continuous PCRs for the insert fragment as described below to construct a gene replacement vector. Primer pairs SGI3-del-F1/SGI3-del-R1 and SGI3-del-F2/SGI3-del-R2 were used to amplify upstream and downstream regions of SGI3, respectively (1^st^ PCR). Target sites of primers SGI3-del-R1 and SGI3-del-F2 are *attL* and *attR*, respectively, and the primer sequence is the complementary each other. First PCR was performed in a 20-µl reaction mixture containing template DNA from L-3841, 0.3 µM each primer, 0.2 mM deoxynucleotide triphosphates (dNTPs), 1 mM MgSO_4_, PCR buffer, and 0.4 U KOD-Plus-DNA polymerase (Toyobo Co., Ltd., Osaka, Japan) using 25 amplification cycles. PCR products were purified using the QIAquick Gel Extraction Kit (QIAGEN, Hilden, Germany). Second PCR was performed in a 30-µl reaction mixture containing the first PCR products, 0.2 mM dNTPs, 1 mM MgSO_4_, PCR buffer, and 0.3 U KOD-Plus-DNA polymerase using 2 amplification cycles. Third PCR was performed in a 50-µl reaction mixture containing 30 µl of the 2^nd^ PCR product as a template, 0.3 µM outermost primers, SGI3-del-F1 and SGI3-del-R2, 0.2 mM dNTPs, 1 mM MgSO_4_, PCR buffer, and 0.5 U KOD-Plus-DNA polymerase using 25 amplification cycles. The same procedure was performed to generate the insert fragments for the gene replacement vector for *pco* gene cluster and *cus* gene cluster. All primers used for the procedure are listed in Table S1. Each fragment was cloned into SmaI-digested temperature-sensitive vector pTH18ks1 (41) in *Escherichia coli* DH5α (Takara Bio Inc.).

The strains L-3841 and LT2TC*pheR* were transformed with one of each vector by electroporation. The cells were spread on LB agar (Becton, Dickinson and Company) supplemented with kanamycin (30 µg/ml) and incubated at 30°C overnight. The colonies were then streaked on the same agar plates pre-warmed at 42°C and incubated at 42°C overnight. The single crossover strains were purified under the same conditions and then passaged at 30°C several times. The double crossover strains were screened for the loss of kanamycin resistance, and the mutation was checked by PCR and sequencing of the amplicon using appropriate primers.

### Heavy metal susceptibility testing

MICs of strains to CuSO_4_, ZnCl_2_, and Na_2_HAsO_4_ (FUJIFILM Wako Pure Chemical) were determined by the agar dilution method using Mueller-Hinton agar plates (Becton, Dickinson and Company). The spotted plates were incubated at 37°C under aerobic conditions for 20 hours or anaerobic conditions (anaerobic jar with AnaeroPack-Anaero; Mitsubishi GAS Chemical Company, Inc., Tokyo, Japan) for 48 hours.

### Accession number(s)

Complete genome sequences of *S. enterica* strains L-3841 and L-3838 were deposited in the DDBJ Sequence Read Archive under accession numbers AP019374 and AP019375, respectively.

## RESULTS

### Structure of SGI3 in wild-type strains

The clade 9 isolates L-3838 and L-3841 harbored a genomic island, which was identical to SGI3 in the Australian *Salmonella* 4,[5],12:i:-strain TW-Stm6 (33). The ORFs of SGI3 in L-3841 were listed in Table S2. The SGI3 length of L-3838 and L-3841 is 80,686 bp except for a direct repeat in both ends, and the nucleotide sequence identity of both isolates was 100%. SGI3 of these isolates had almost identical nucleotide sequence (80,685/80,686 bp) to that of TW-Stm6. SGI3 of L-3838 was integrated into the 3′ end of the chromosomal tRNA gene *pheV* and formed a 52-bp direct repeat at both ends. SGI3 of L-3841 was integrated into the 3′ end of the chromosomal tRNA gene *pheR* and formed a 55-bp direct repeat at both ends (Fig. S1). Among eighty-six ORFs identified in L-3841, 24 ORFs were predicted as heavy metal-resistant genes (Fig. 1). Seventeen out of the 24 ORFs formed a heavy metal homeostasis/resistance island called the Copper Homeostasis and Silver Resistance Island (CHASRI) (42). The CHASRI consists of a plasmid-borne copper resistance system (*pco*) cluster (*pcoABCDRSE*) (43), a *sil* heavy metal export system cluster (*silE* and *silP*) (44), and a copper sensing copper efflux system (*cus*) cluster (*cusABFCRS*) (45). The nucleotide sequence identity of CHASRI located in SGI3 and *E. coli* str. APEC-O1 plasmid pAPEC-O1-R (GenBank accession number NC_009838.1) was 95% (Fig. 2). The remaining seven heavy metal-resistant genes formed an arsenic resistance operon (*arsRDABC*) (46). The nucleotide sequence identity of the arsenic operon located in SGI3 and *Enterobacter cloacae* ATCC13047 plasmid pECL_A (GenBank accession number CP001919.1) was 92% (Fig. 2). The structure of the remaining part of SGI3, including genes relating to conjugative transfer and DNA replication or partitioning, showed high similarity with the ICE-like genomic island of *Edwardsiella icutaluri* MS-17-456 (GenBank accession number CP028813.1). The nucleotide sequence identity of genes *traD, traG*, and *traI* located in SGI3 and ICE-like genomic island of *E. icutaluri* MS-17-456 were 91%, 92%, and 86%, respectively (Fig. 2).

**FIG 1.**
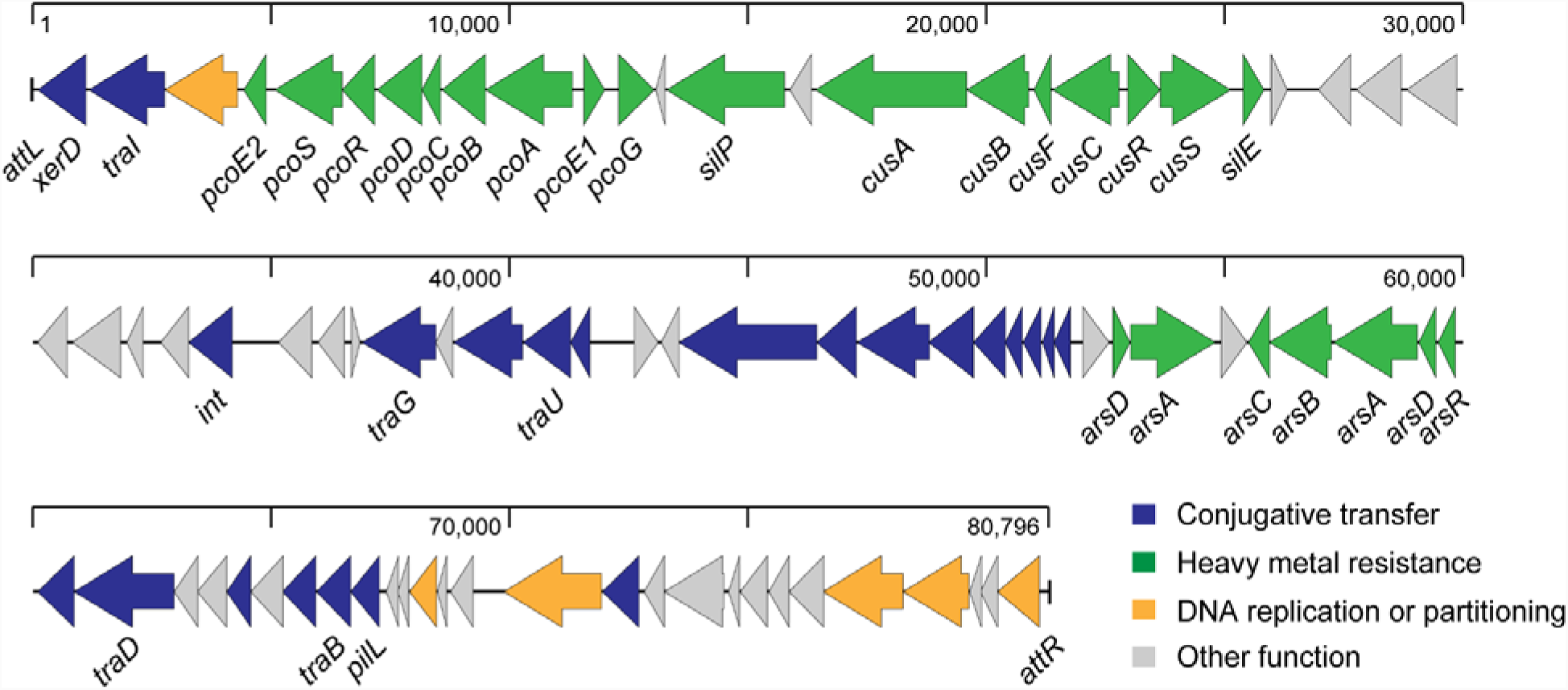
Schematic view of full-length SGI3 of *Salmonella* 4,[5],12:i:-L-3841. The diagram shows the predicted classification of each gene, which is represented as arrows according to the following scheme: violet, genes for conjugative transfer; green, genes for heavy metal resistance; orange, genes for DNA replication or partitioning; gray, genes with other functions.

**FIG 2.**
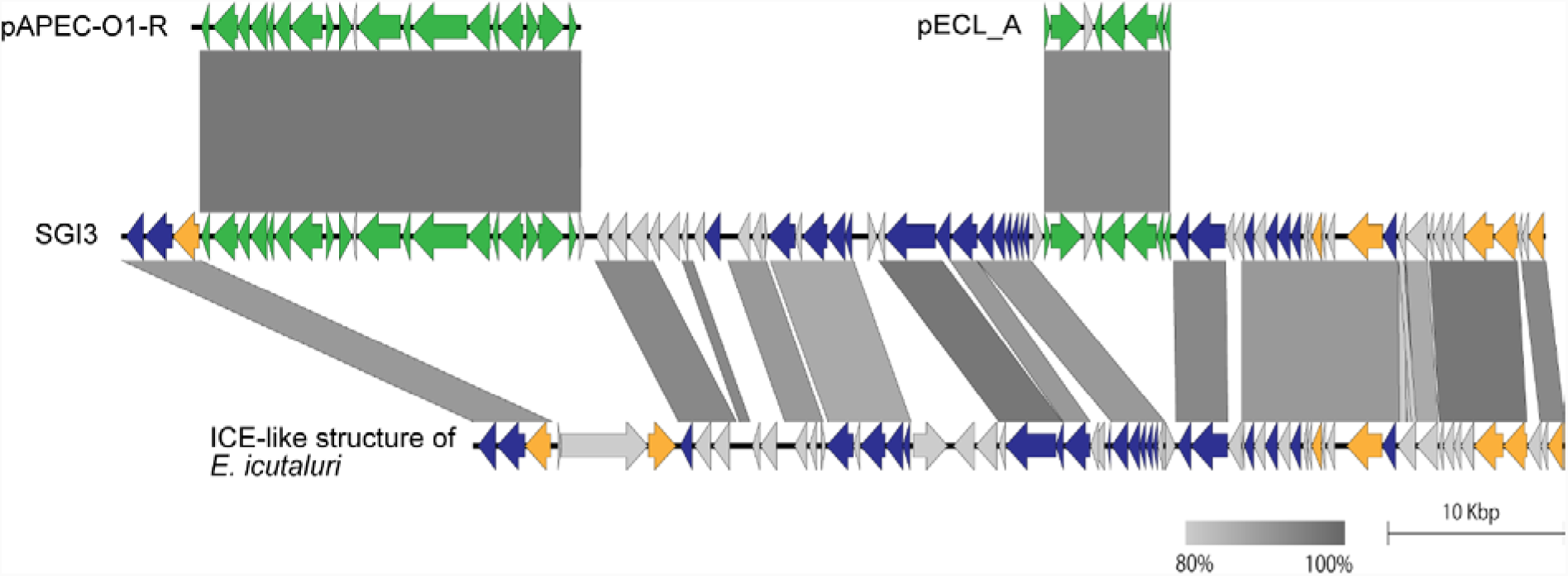
Pairwise alignment of SGI3 of *Salmonella* 4,[5],12:i:-L-3841 with the relevant mobile genetic elements, pAPEC-01-R of *Escherichia coli* APEC-O1, pECL_A of *Enterobacter cloacae* ATCC13047, and ICE-like chromosomal island of *Edwardsiella icutaluri* MS-17-456.

### Transfer of SGI3

The results of the conjugation experiment are shown in Table 2. SGI3 of donor strain *Salmonella* 4,[5],12:i:-L-3841 (SNP genotype 9) was successfully transferred to recipient strains of *Salmonella* Typhimurium or *Salmonella* 4,[5],12:i:-with SNP genotypes 1 to 8. The transfer frequencies ranged from 1.2 × 10^−7^ to 1.3 × 10^−4^ per donor. The SGI3 was also transferred to other serovars, including Heidelberg, Thompson, Hadar, Newport, and Cerro, with the transfer frequencies ranging from 4.1× 10^−9^ to 1.7 × 10^−5^ per donor.

**Table 2.** Transfer frequency of SGI3 to *S. enterica* strains

| Strain | Recipient |  |  | Transfer frequency<sup>b</sup><br>(per donor) |
| --- | --- | --- | --- | --- |
|  | Serovar | O group | SNP genotype<sup>a</sup> |  |
| L-3900 | Typhimurium | 4 | 1 | $1.4 \times 10^{-6}$ |
| L-4131 | Typhimurium | 4 | 2 | $7.0 \times 10^{-5}$ |
| LT2 | Typhimurium | 4 | 3 | $1.2 \times 10^{-7}$ |
| L-3785 | 4,[5],12:i:- | 4 | 4 | $4.5 \times 10^{-5}$ |
| L-4153 | Typhimurium | 4 | 5 | $1.3 \times 10^{-4}$ |
| L-4125 | Typhimurium | 4 | 6 | $2.3 \times 10^{-5}$ |
| L-4239 | Typhimurium | 4 | 7 | $7.3 \times 10^{-5}$ |
| L-3569 | Typhimurium | 4 | 8 | $2.4 \times 10^{-5}$ |
| L-2707 | Heidelberg | 4 | NA | $1.7 \times 10^{-5}$ |
| L-3038 | Infantis | 7 | NA | ND |
| L-3202 | Choleraesuis | 7 | NA | ND |
| L-4000 | Thompson | 7 | NA | $6.1 \times 10^{-8}$ |
| L-2819 | Hadar | 8 | NA | $4.1 \times 10^{-9}$ |
| L-3037 | Newport | 8 | NA | $7.4 \times 10^{-6}$ |
| L-3046 | Dublin | 9 | NA | ND |
| L-3241 | Enteritidis | 9 | NA | ND |
| L-3289 | Cerro | 18 | NA | $6.4 \times 10^{-8}$ |

<sup>a</sup>NA, not applicable<sup>b</sup>ND, not detected

### Localization of SGI3 in the recipients

To confirm the chromosomal location of SGI3 among the donor and transconjugants, PCR, direct sequencing of the products, and PFGE-Southern blot analysis were performed. As shown in Fig. 3a, strains LT2, LT2TC*pheV*, and LT2TC*pheR* were identified as SNP genotype 3, whereas L-3838 and L-3841 were identified as SNP genotype 9 by the SNP genotyping system. PCR and the direct sequencing of the products was confirmed that the SGI3 insertion site of strains L-3838 and LT2TC*pheV* was the 3′ end of *pheV*, whereas that of L-3841 and LT2TC*pheR* was the 3′ end of *pheR*. The *attI* sequence indicating the generation of circular form of SGI3 was also detected from wild types and recipients by PCR and direct sequencing of the products.

**FIG 3.**
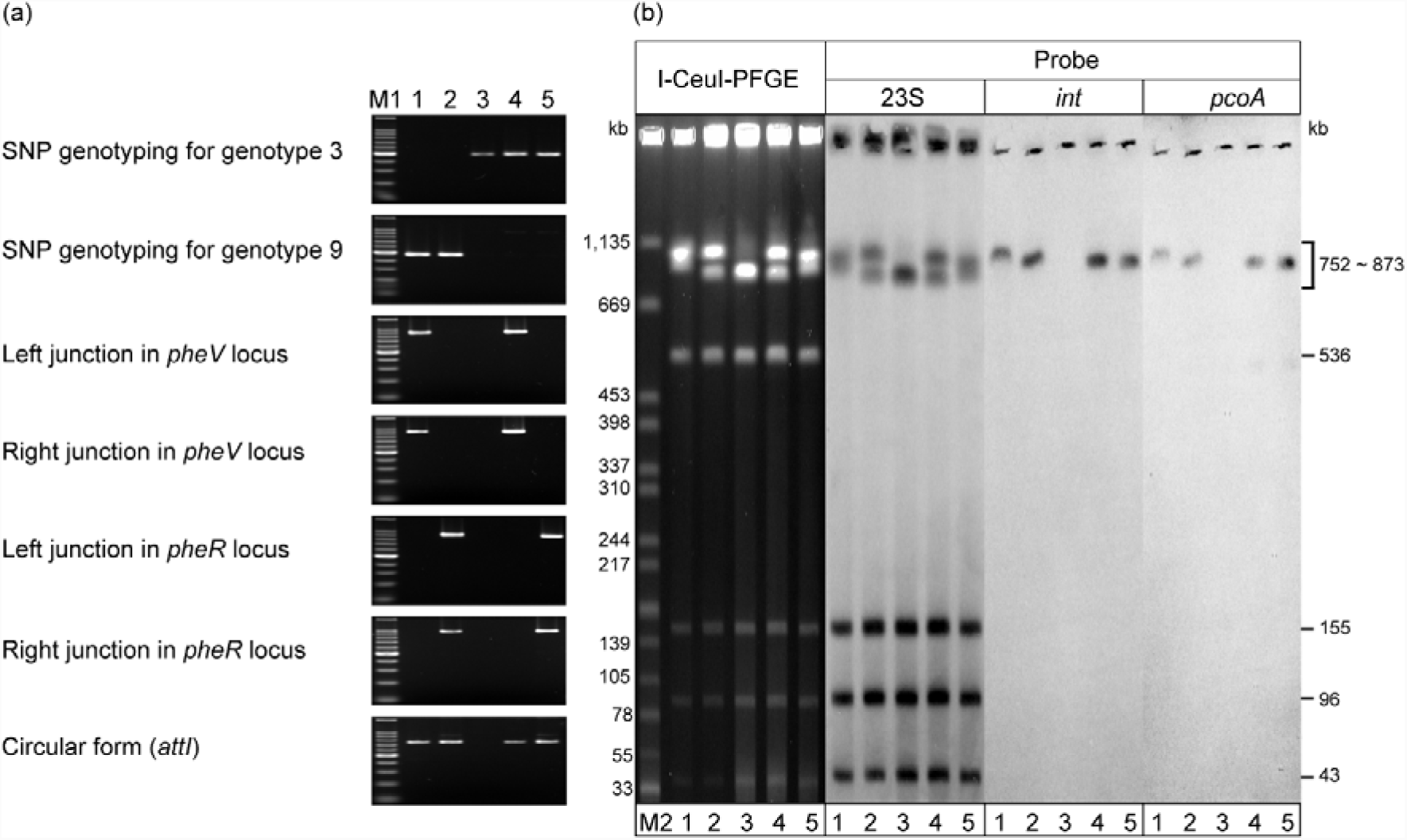
PCR and PFGE-Southern blot hybridization images demonstrating the chromosomal location and/or generation of the circular form of SGI3 in *Salmonella* 4,[5],12:i:-strains and *Salmonella* Typhimurium LT2 transconjugants. Lane 1, L-3838; lane 2, L-3841; lane 3, LT2; lane 4, LT2TC*pheV*; lane 5, LT2TC*pheR*; lane M1, 100-bp DNA ladder; lane M2, XbaI-digested *Salmonella* Braenderup. (a) SNP genotyping and PCR to detect junction regions of integrated or circular form of SGI3. (b) PFGE separation of I-CeuI-digested genomic DNA from *S. enterica* strains followed by Southern blot hybridization with 23S rRNA, integrase, and *pcoA* gene probes.

LT2 contains seven I-CeuI restriction sites located on the 23S rRNA genes on the chromosome (GenBank accession number NC_003197.1). I-CeuI digestion of the LT2 chromosome generated fragments 2,504, 770, 752, 536, 155, 96, and 43 kb in size. As shown in Fig. 3b, the 2,504-kb fragment did not migrate from the well, and the 770-kb and 752-kb fragments were indistinguishable by the PFGE conditions used in this study. To detect the chromosomal fragments by Southern blot analysis, the probe 23S, which targeted the 23S rRNA gene sequence, was used. This probe hybridized with six fragments 2504, 770, 752, 155, 96, and 43 kb in size, as expected. The expected SGI3 insertion sites *pheV* and *pheR* are located on the 770-kb and 752-kb fragments, respectively. Thus, when SGI3 was inserted into the *pheV* and *pheR* locus in LT2, 851-kb and 833-kb fragments should be generated in the transconjugants instead of 770-kb and 752-kb fragments, respectively. The expected fragments were identified in strains LT2TC*pheV* and LT2TC*pheR*, respectively. Sizes of SGI3 containing I-CeuI-digested fragment of strains L-3838 and L-3841 were expected as 861-kb and 873-kb, respectively. Both *int* and *pcoA* gene probes hybridized with the 861-kb fragment of L-3838, 873-kb fragment of L-3841, 852-kb fragment in LT2TC*pheV*, and 833-kb fragment in LT2TC*pheR*, which were generated from the chromosome. Direct sequencing of the amplified fragments of the junction regions identified direct repeats of 52 and 55 bp in strains LT2TC*pheV* and LT2TC*pheR*, respectively, as shown in Fig. S1. The size of the *attI* sequence of the circular form of SGI3 was 55 bp in the both strains.

### Susceptibility of wild types, **transconjugants**, **and deletion mutants to heavy metals**

The MICs of the SNP genotype 9 strains to CuSO_4_ under anaerobic condition were four to six times higher than those of other SNP genotypes of *Salmonella* Typhimurium, serovars Heidelberg, Hadar, Newport, Cerro, Thompson, Infantis, and Enteritidis. Serovars Dublin and Choleraesuis were slightly more susceptible to CuSO_4_ under anaerobic conditions. The MICs of the wild-type strains and their transconjugants to CuSO_4_ were the same under aerobic conditions with the exception of strain L-4153. The MICs of transconjugants to CuSO_4_ under anaerobic conditions were four to six times higher than those of their wild-type strains. The MICs were the same as that of the donor strain L-3841. The MICs of wild-type strains and their transconjugants to ZnCl_2_ were the same or slightly different (1 mM difference) under both aerobic and anaerobic conditions. The MIC of genotype 9 strains to Na_2_HAsO_4_ under aerobic conditions was >64, whereas that of other genotypes of *Salmonella* Typhimurium and other serovars were ranged from < 0.25 to 4. The MICs of the transconjugants to Na_2_HAsO_4_ were at least 32-fold increased compared with those the wild-type strains. The MICs of LT2TC*pheR*ΔSGI3 and LT2TC*pheR*Δ*cus* to CuSO_4_ under anaerobic conditions were six times lower than those of their parental strains, LT2TC*pheR* (Table 3).

**Table 3.** Susceptibility of strains to heavy metals

| Strain | MIC (mM) |  |  |  |  |
| --- | --- | --- | --- | --- | --- |
|  | CuSO<sub>4</sub> |  | ZnCl<sub>2</sub> |  | Na<sub>2</sub>HAsO<sub>4</sub> |
|  | Aerobic | Anaerobic | Aerobic | Anaerobic | Aerobic |
| L-3838 | 8 | 6 | 2 | 3 | > 64 |
| L-3841 | 8 | 6 | 3 | 2 | > 64 |
| L-3841ΔSGI3 | 6 | 1 | 3 | 3 | ≤ 0.25 |
| L-3841Δ*pco* | 8 | 6 | 3 | 3 | > 64 |
| L-3841Δ*cus* | 8 | 1 | 3 | 3 | > 64 |
| L-3900 | 8 | 1 | 3 | 6 | 0.5 |
| L-3900TC | 8 | 6 | 3 | 6 | 64 |
| L-4131 | 8 | 1 | 3 | 3 | ≤ 0.25 |
| L-4131TC | 8 | 6 | 3 | 4 | > 64 |
| LT2 | 8 | 1 | 4 | 6 | 0.5 |
| LT2TC*pheV* | 8 | 6 | 3 | 6 | 64 |
| LT2TC*pheR* | 8 | 6 | 3 | 6 | > 64 |
| LT2TC*pheR* ΔSGI3 | 8 | 1 | 4 | 6 | 0.5 |
| LT2TC*pheR* Δ*pco* | 8 | 6 | 4 | 6 | > 64 |
| LT2TC*pheR* Δ*cus* | 8 | 1 | 4 | 6 | > 64 |
| L-3785 | 8 | 1 | 3 | 3 | 0.5 |
| L-3785TC | 8 | 6 | 3 | 4 | > 64 |
| L-4153 | 8 | 1 | 4 | 3 | 0.5 |
| L-4153TC | 6 | 6 | 3 | 3 | > 64 |
| L-4125 | 8 | 1.5 | 2 | 2 | 0.5 |
| L-4125TC | 8 | 6 | 2 | 2 | > 64 |
| L-4239 | 8 | 1 | 3 | 4 | 0.5 |
| L-4239TC | 8 | 6 | 3 | 4 | 64 |
| L-3569 | 8 | 1 | 4 | 6 | 0.5 |
| L-3569TC | 8 | 6 | 3 | 6 | 64 |
| L-2707 | 8 | 1 | 3 | 3 | ≤ 0.25 |
| L-2707TC | 8 | 6 | 3 | 3 | 64 |
| L-2819 | 6 | 1.5 | 3 | 3 | ≤ 0.25 |
| L-2819TC | 6 | 6 | 3 | 3 | >64 |
| L-3037 | 6 | 1 | 3 | 4 | ≤ 0.25 |
| L-3037TC | 6 | 6 | 3 | 4 | >64 |
| L-3289 | 8 | 1 | 2 | 6 | ≤ 0.25 |
| L-3289TC | 8 | 6 | 2 | 6 | >64 |
| L-4000 | 8 | 1 | 4 | 3 | 4 |
| L-4000TC | 8 | 6 | 4 | 3 | >64 |
| L-3038 | 6 | 1 | 3 | 3 | ≤ 0.25 |
| L-3046 | 8 | 0.75 | 3 | 4 | 0.5 |
| L-3202 | 3 | ≤ 0.5 | 3 | 4 | ≤ 0.25 |
| L-3241 | 8 | 1 | 3 | 3 | 0.5 |

## DISCUSSION

In this study, we characterized the genomic island SGI3, which was first described in the European *Salmonella* 4,[5],12:i:-clone (15). The corresponded clone has been isolated in Japan since 2000s and was designated as clade 9 (SNP genotype 9) in the previous study (12). The SGI3 structure of Japanese strains was almost identical to that of the Australian strain TW-Stm6 (33) and the European strain SO4698-09 (15). The SGI3 nucleotide sequence identity of these isolates was almost 100% across the entire length of 80,685 bp. These data support the notion that the worldwide dissemination of the *Salmonella* 4,[5],12:i:-clone originated from Europe.

The nucleotide sequences of the core region of SGI3 showed low similarity to representative ICEs that were reported previously. However, SGI3 contains genes encoding a site-specific recombinase XerD (47), an ICE relaxtase TraI (48), and typeVI secretion system (Table S2), suggesting that this island is a mobile genetic element. Transconjugants that acquired the SGI3 in their chromosome were successfully selected after the filter mating experiment in this study (Table 2). The 3′ end of tRNA genes is a representative attachment site of ICEs reported previously (49), and SGI3 was inserted into the 3′ end of tRNA genes *pheV* or *pheR* (Fig. 3a). Southern blot analysis results supported the chromosomal location of SGI3 in the transconjugants (Fig. 3b). The *attI* sequence indicating the generation of circular form of SGI3 was detected by PCR in donor and recipient strains (Fig. 3a). These data suggested that SGI3 was excised from the donor chromosome, transferred by conjugation, and integrated into the host chromosome by site-specific recombination.

A direct repeat (*attL* and *attR*) was formed by the integration of SGI3 in the chromosome of the transconjugants. Although the size of the direct repeat generated from *pheV* (52 bp) was 3 bp shorter than that from *pheR* (55 bp), the nucleotide sequence was identical (Fig. S1). The 52-bp and 55-bp sequences seemed to be recognized as attachment sites of SGI3.

We could not detect any transconjugants from four strains of *S. enterica*, including serovars Infantis, Choleraesuis, Dublin, and Enteritidis, after three independent conjugation experiments, while transconjugants were successfully obtained from all genotypes of *Salmonella* Typhimurium, including *Salmonella* 4,[5],12:i:-. We detected the 55-bp attachment sequences among these strains by PCR and sequencing (data not shown). In this study, we performed the conjugation experiment without any chemicals inducing the SOS response. The addition of mitomycin C or antimicrobials might leads to successful conjugative transfer of SGI3 to these strains (19).

SGI3 contains CHASRI and the *ars* gene cluster, which contribute to heavy metal-resistance (Fig. 1). CHASRI emerged and evolved in response to copper deposition across aerobic and anaerobic environments (42). CHASRI confers copper resistance under aerobic and anaerobic conditions and during the shift between aerobic and anaerobic environments, which could explain its wider distribution among facultative anaerobes.

CHASRI consists of two copper-resistance gene clusters: *pco* and *cus*. The *pco* gene cluster includes *pcoABCDE1E2G-RS* and is potentially involved in periplasmic copper detoxification (50). PcoA is a multi-copper oxidase and oxidizes Cu(I) to less toxic Cu(II) in the periplasm. Since PcoA has a twin-arginine motif in its leader sequence, it is probably translocated into the periplasm via the twin-arginine translocation (TAT) pathway with copper bound to its active sites. To load copper into catalytic sites of PcoA, PcoC and PcoD might transport copper across the cytoplasmic membrane. PcoE1 and PcoE2 bind copper in the periplasm and possibly carry copper to PcoA. The *cus* gene cluster consists of six genes: *cusCFBA-RS* (51). On the other hand, CusABC is a tripartite copper-efflux pump consisting of an inner membrane pump (CusA), a periplasmic protein (CusB), and an outer membrane protein (CusC) and forms a channel spanning the entire cell envelope (50). CusF is a periplasmic metallochaperone that can bind Cu(I) or Ag(I) and transfer those to CusB metal-binding sites. The metal-bound CusB is required for activation of the metal transfer to a binding site of the CusA antiporter (52). PcoRS and CusRS are copper-response two-component regulatory systems (50), and the latter has an important role for copper resistance under anaerobic conditions in *E. coli* (53).

*Salmonella* Typhimurium intrinsically possesses a CueR regulon composed of genes encoding a copper-sensing transcriptional regulator CueR, copper efflux pump CopA, and a multicopper oxidase CueO to maintain intracellular copper homeostasis (54, 55). It is reported that CueR and CusRS sensor systems are differentially regulated in response to changes in oxygen availability (53). Expression of *cueO* and *copA* under aerobic conditions was higher than that under anaerobic conditions, whereas expression of *cusC* under anaerobic conditions was higher than that under aerobic conditions. These observations suggest that the products of the *cus* gene cluster play an important role in copper homeostasis especially under anaerobic conditions.

In our study, all of the SGI3 transconjugants had higher MICs to CuSO_4_ compared with the recipient strains under anaerobic conditions. Furthermore, the MICs of the *cus* gene cluster deletion mutants L-3841Δ*cus* and LT2TC*pheR*Δ*cus* to CuSO4 were equivalent to those of L-3841ΔSGI3 and LT2TC*pheR*ΔSGI3, respectively (Table 3). These data indicate that the products of the *cus* gene cluster contribute to copper resistance of *S. enterica* under anaerobic conditions and are concordant with results of earlier studies (53-55). On the other hand, we did not detect MIC differences between *pco* gene cluster deletion mutants and parental strains. Given that the contribution of the *pco* gene cluster products to copper resistance is subtle (57), intrinsic copper resistance under aerobic conditions might mask this activity.

The *ars* gene cluster includes *arsABCDR* (58). ArsA is a soluble ATPase, which interacts with the integral membrane protein ArsB. Prior to efflux, arsenate is reduced to arsenite by small cytoplasmic arsenate reductase ArsC to activate ATPase activity of ArsA. Then, arsenite is pumped out through ArsB. This system involves the detoxification of As(III) and AS(V). All of the transconjugants obtained in this study had distinctly higher MICs to Na_2_HAsO_4_ compared with recipient strains under aerobic conditions. Deletion of SGI3 in the strains L3841 and LT2TC*pheR* reduced remarkably their MICs, while deletion of the gene clusters *pco* and *cus* did not affect their MICs. These data suggest that the *ars* gene cluster contributes to arsenic resistance of *S. enterica* under aerobic conditions.

Heavy metals, such as copper and zinc, are added to livestock feed as micronutrients for growth promotion and suppressing gut pathogens (16). Aromatic organoarsenic compounds are also used as feed additives in the poultry industry for the same purposes (59). Medardus *et. al*. (2014) reported that copper levels in swine feed samples ranged between 3.2 and 365.2 mg/kg. Copper levels in fecal samples were higher, ranging from 71.2 to 2,397 mg/kg. Zinc levels in swine feed samples varied between 77 and 2,000 mg/kg. The level of zinc in fecal samples was significantly higher, ranging between 536.5 and 12,557.2 mg/kg (16). The use of high concentrations of heavy metals as feed additives and biocides and the subsequent deposition across the environment might pose strong selection pressures to bacteria. Notably, SGI3 conferred copper resistance under anaerobic conditions, suggesting that the acquisition of SGI3 might support *S. enterica* growth in the intestinal environment containing higher concentrations of heavy metals.

In conclusion, the findings of this study clearly demonstrate that SGI3 is an ICE and elevates host resistance to copper and arsenic. The European clone has been one of the major clones of *Salmonella* Typhimurium worldwide for more than 20 years. Elevated usage of heavy metals as feed additives might represent a selection pressure for this clone. Continuous monitoring of heavy metals as feed additives and the resistance of bacteria detected in the livestock industry is required.

## ACKNOWLEDGMENTS

The authors are grateful to the staff at the prefectural livestock hygiene service centers for providing *Salmonella* isolates. This study was supported by grant-in-aid for the Research Program on Emerging and Reemerging Infectious Diseases (JP18fk0108048) from the Japan Agency for Medical Research and Development (AMED).

